# Prevalence study of cellular capsid-specific immune responses to AAV2, 4, 5, 8, 9 and rh10 in healthy donors

**DOI:** 10.1101/2023.12.20.570598

**Authors:** Rebecca Xicluna, Allan Avenel, Céline Vandamme, Marie Devaux, Nicolas Jaulin, Célia Couzinie, Johanne Le Duff, Alicia Charrier, Mickaël Guilbaud, Oumeya Adjali, Gwladys Gernoux

**Affiliations:** Nantes Université, CHU de Nantes, INSERM, TaRGeT – Translational Research in Gene Therapy, UMR 1089, F-44200 Nantes, France

**Keywords:** AAV, gene therapy, prevalence, cellular immune response, anti-AAV T cells and anti-AAV IgG

## Abstract

Recombinant adeno-associated virus (rAAV) vectors appear, more than ever, to be efficient viral vectors for *in vivo* gene transfer as illustrated by the approvals of 7 drugs across Europe and the USA. Nevertheless, pre-existing immunity to AAV capsid in humans remains one of the major limits for a successful clinical translation. Whereas pre-existing humoral response to AAV capsid is well documented, the prevalence of pre-existing capsid-specific T cell responses still needs to be studied and characterized. Here, we investigated the prevalence of AAV-specific circulating T cells towards AAV2, 4, 5, 8, 9 and rh10 in a large cohort of healthy donors using the standard IFNγ ELISpot assay. We observed the highest prevalence of pre-existing cellular immunity to AAV9 serotype followed by AAV8, AAV4, AAV2, AAVrh10 and AAV5 independently of the donors’ serological status. An in-depth analysis of T cell responses towards the 2 most prevalent serotypes 8 and 9 shows that IFNγ secretion is mainly mediated by CD8 T cells for both serotypes. A polyfunctional analysis reveals different cytokine profiles between AAV8 and AAV9. Surprisingly, no IL-2 secretion was mediated by anti-AAV9 immune cells suggesting that these cells may rather be exhausted or terminally differentiated than cytotoxic T cells. Altogether, these results suggest that pre-existing immunity to AAV may vary depending on the serotype and support the necessity of using multiparametric monitoring methods to better characterize anti-capsid cellular immunity and foresee its impact in rAAV-mediated clinical trials.

## INTRODUCTION

Recombinant adeno-associated viral vectors (rAAV)-mediated gene therapy shows promising benefits for the treatment of genetic disorders as illustrated by Glybera®, Luxturna®, Zolgensma®, Upstaza®, Roctavian®, Hemgenix® and Elevidys® FDA and EMA approvals and more than 230 clinical trials ongoing (^1^ and clinicaltrials.gov). However, there are still major hurdles to overcome for their successful clinical translation such as their immunogenicity in patients. Indeed, since protocols evolved from local to systemic delivery with high doses of vector, adverse events related to the immune system activation have been reported in treated patients, resulting in some cases in regulatory holds of clinical trials.^2^

Seroprevalence studies showed that human population is exposed to wild-type (WT) AAV virus early in life^3^ and might present a memory immune response that can be reactivated after gene transfer. These studies were mainly conducted on human serum samples to evidence the presence of anti-AAV neutralizing antibodies^4,5^ that will prevent target cell transduction after vector administration but little is known about the anti-AAV memory T cell response prevalence in humans. Importantly, these pre-existing anti-AAV memory T cells might be reactivated after gene transfer and lead to gene transfer failure as previously described in a clinical trial targeting Hemophilia B patients.^6,7^ In this trial, the intrahepatic injection of rAAV2 led to the destruction of liver transduced cells 4-8 weeks after vector delivery due to the reactivation of pre-existing IFNγ-secreting AAV-specific CD8 T cells. Since then, the liver transaminase elevation due to hepatotoxicity and the IFNγ secretion in response to AAV capsid are systematically measured in clinical trials and managed by corticosteroid treatment.^8–10^

To our knowledge, in the general population, only cellular responses directed to AAV2, AAV1, AAV5 and AAV8^11–13^ have been investigated so far, which does not cover the whole diversity of rAAV serotypes now used in clinical trials. Indeed, no cellular prevalence data in humans are reported for the AAV9 capsid which is used in a commercialized product since 2019 for the treatment of spinal muscular atrophy (SMA) in pediatric patients.^14^ AAV9 serotype is also used in clinical trials targeting Pompe (NCT02240407), Canavan (NCT04998396), GM1 gangliosidosis (NCT03952637) and Parkinson (NCT04127578) diseases as well as Duchenne Muscular Dystrophy (NCT04240314, NCT03368742 and NCT03362502). The AAVrh10 has also emerged lately as a good candidate for gene delivery to the central nervous system and showed promising results for clinical applications targeting Krabbe disease,^15^ metachromatic leukodystrophy (^16^ and NCT01801709), GM1 gangliosidosis (^17^ and NCT04273269) or amyotrophic lateral sclerosis (ALS).^8,18^

Here, we investigated the prevalence of AAV-specific circulating T cells towards AAV2, 4, 5, 8, 9 and rh10 in a large cohort of healthy donors using an IFNγ ELISpot assay that is usually used in immune monitoring in clinical studies. We observed the highest prevalence of pre-existing cellular immunity against AAV9 serotype followed by AAV8, AAV4, AAV2, AAVrh10 and AAV5. As already described, there was no correlation in our cohort between the cellular immune response and the humoral anti-capsid immunity. An in-depth analysis of T cell responses towards the 2 most prevalent serotypes 8 and 9 shows that IFNγ secretion is mainly mediated by CD8 T cells for both serotypes. We also performed FluoroSpot assays that showed their potential as a polyfunctional assay to characterize anti-AAV T cell responses to be used in clinical trials as we observed different cytokine profiles between AAV8 and AAV9. This observation suggests a different functional impact of preexisting capsid-specific T cells in rAAV-mediated clinical trials.

Our data suggest that pre-existing anti-AAV cellular immune responses are serotype-dependent and encourage the use of multiparametric monitoring methods to better characterize capsid-specific cellular immune responses and predict their impact in rAAV-mediated clinical trials.

## MATERIALS AND METHODS

All this work was performed at INSERM TaRGeT UM1089, Nantes, France under the control of our quality management system that is approved by Lloyd’s Register Quality Assurance LRQA to meet requirements of international Management System Standards ISO 9001:2015. It has been implemented to cover all activities in the laboratory, including research experiments and production of research-grade viral vectors.

### Healthy donor blood samples

Cytapheresis samples were provided by the local Etablissement Français du Sang (EFS Nantes, Pays de la Loire, agreements N° PLER NTS 2016-25 and CPDL-PLER-2021 005) and originated from consenting healthy donors living in the Pays de la Loire area, France (22 women and 129 men with age ranging from 19 to 72 years old). All the participants gave their written informed consent. PBMCs were isolated using Ficoll density gradient centrifugation (Ficoll-Paque PLUS, GE Healthcare) and frozen in liquid nitrogen for ELISpot and Fluoropost assays. Plasma were isolated by spinning 10min at 2000rpm, RT 10ml of whole blood and were stored at -20°C.

### IFNγ ELISpot assay on healthy donor PBMCs

Anti-AAV cellular immune responses were analyzed using an IFNγ ELISpot assay. ELISpot assays were performed according to the manufacturer’s recommendations (ELISpot Plus: Human IFN-y (ALP) kit, MABTech). Briefly, PBMCs from healthy donors were stimulated *in vitro* for 18-48h with overlapping peptides spanning the VP1 capsid protein sequence (which also covers VP2 and VP3) of each serotype (*i.e.* AAV2, 4, 5, 8, 9 and rh10; NCBI accession numbers AAC03780, AAC58045.1, AAD13756.1, AAN03857.1, AAS99264.1 and AAO88201.1 respectively) and divided into 3 pools (15-mers overlapping by 10 aa, Sigma-Aldrich, United States or Mimotopes, Australia). For each serotype, 143 (AAV5), 145 (AAV2 and 4) or 146 (AAV8, 9 and rh10) peptides were synthetized. Pool 1 was made of peptides 1-46 (AAV2), 1-48 (AAV5) or 1-49 (AAV4, 8, 9 and rh10). Pool 2 was made of peptides 47-96 (AAV2), 49-96 (AAV5) or 50-98 (AAV4, 8 and 9) or 50-97 (AAVrh10). Pool 3 was made of peptides 97-145 (AAV2), 97-143 (AAV5), 99-146 (AAV4, 8 and 9) or 98-146 (AAVrh10). A negative control consisted of unstimulated cells (medium only), whereas concanavalin A (Con A, 10 µg/mL, Sigma-Aldrich, United States) stimulation was used as a positive control for cytokine secretion. Spot number was determined using an ELISpot iSpot Spectrum reader (AID, Strassberg, Germany) and a Mabtech IRIS reader and analyzed with AID ELISpot reader Software V7.0 (AID, Strassberg, Germany) and Mabtech APEX v1.1 software (Mabtech, Sweden) respectively. Responses were considered positive when the number of spot-forming colonies (SFCs) *per* 1e6 cells were >50 and at least 3-fold higher than the negative control condition.

### AAV vector productions

For ELISA assays, single-stranded AAV serotypes 2, 8, 9 and rh10 vectors were produced by the Center for Production of Vectors (CPV-vector core from University Hospital of Nantes/French Institute of Health [INSERM], University of Nantes [https://umr1089.univ-nantes.fr/en/facilities-cores/cpv]). Briefly, vectors were produced by cotransfection of human embryonic kidney (HEK) 293 cells with the vector plasmid and helper plasmid (containing helper genes from adenovirus and the *rep cap* genes according to the capsid serotype). Vectors were purified by cesium-chloride gradient. The enriched-empty (AAV2, 9 and rh10) or enriched-full (AAV8) particles band were collected and dialyzed. The amount of viral capsid proteins for the AAV2, AAV9 and AAVrh10 vector batches were compared to an AAV2 standard by quantitative SDS-PAGE Coomassie blue staining. An ELISA total capsid titration was performed for the AAV8 (kit from Progen, Ref. #PRAAV8).

### Anti-AAV IgG ELISA

Detection of anti-AAV total IgG antibodies for each serotype in the plasma of healthy human donors was conducted using an enzyme-linked immunosorbent assay (ELISA). Nunc Maxisorp P96 plates (Sigma-Aldrich) were coated overnight at 4°C with rAAV2, rAAV8, rAAV9 or rAAVrh10 particles (INSERM UMR 1089 - Center for Production of Vectors). Vectors were produced, purified and titrated as described in the previous section. After washing and plate saturation, wells were then incubated for 1 hour at 37°C with at least 6 serial 4-fold dilutions of human plasma starting at 1:10 followed by HRP-conjugated anti-human F(ab’)2 IgG antibody (Jackson) for 1 hour at 37°C. Revelation was performed using 3,3’-5,5’-Tetramethylbenzidine (TMB, OptEIA, BD Biosciences) and absorbance of duplicate samples was read at 450 nm with a correction at 570 nm on a MultiSkan Go reader (Thermo Scientific). Positive threshold curves for each ELISA were determined from seronegative human sera (> 20 donors) as the mean optic density (O.D.) for each dilution + 2*SD. For each donor, anti-AAV IgG titer was defined as the last plasma dilution with an O.D. remaining above the threshold curve.

### IFNγ ELISpot assay on healthy donor CD4- or CD8-depleted PBMCs

CD4+ or CD8+ T cells were depleted from total PBMCs using Stem Cell magnetic-based system (EasySep^TM^ Human CD4/CD8 Positive Selection Kit II) according to manufacturer’s recommendations. Cell depletion efficiency was determined by flow cytometry before (total PBMCs) and after magnetic separation (CD4- and CD8-depleted PBMCs). Briefly, CD4+ T cell and CD8+ T cell populations were analyzed using CD3 (557749; BD Biosciences), CD19 (561031; BD Biosciences), CD4 (560251; BD Biosciences) and CD8 (93-0088-42; eBiosciences) markers. CD4 and CD8 T cells were gated as CD4+ or CD8+ cells respectively within the CD3+CD19-population. Cells were acquired using a BD FACS LSRII flow cytometer (BD Biosciences) and analyzed with FlowJo^TM^ software (v.10; BD Life Sciences). Anti-AAV cellular responses were assessed using the IFNγ ELISpot assay described above.

### IFNγ/IL-2/TNFα Fluorospot assay on healthy donor PBMCs

Anti-AAV cellular immune responses were also evaluated using IFNγ/IL-2/TNFα FluoroSpot assays. FluoroSpot assays were performed according to the manufacturer’s recommendations (FluoroSpot Plus: Human IFNγ/IL-2/TNFα kit, MABTech). Briefly, PBMCs from healthy donors were stimulated *in vitro* for 48h with the same AAV8 and AAV9 peptide libraries used for IFNγ-ELISpot assays described above. A negative control consisted of unstimulated cells (medium only), whereas concanavalin A (Con A, 10 µg/mL, Sigma-Aldrich, United States) stimulation was used as a positive control for cytokine secretion. Spot number was determined using an ELISpot iSpot Spectrum reader (AID, Strassberg, Germany) and analyzed with AID ELISpot reader Software V7.0 (AID, Germany). Responses were considered positive when the number of spot-forming colonies per million cells was > 50 and at least threefold higher than the medium alone negative control.

## RESULTS

### Cellular immune response prevalence to AAV2, 4, 5, 8, 9 and rh10 in healthy donors

The prevalence of preexisting cellular immune response to AAV serotypes 2, 4, 5, 8, 9 and rh10 was determined using an IFNγ ELISpot assay. Peripheral blood mononuclear cells (PBMCs) were isolated from whole blood collected on healthy human donors (n=45-145 depending on serotypes) from the Pays de la Loire area, France. These cells were restimulated *in vitro* with an overlapping peptide library spanning the VP1 sequence of each serotypes tested (*i.e.* AAV2, 4, 5, 8, 9 and rh10) and divided into 3 pools. Our results show that AAV9 and AAV8 have the highest prevalence: 46% (n=66/145) and 24% (n=35/145) of positive donors respectively followed by AAV4 (10%, n=8/80), AAVrh10 (9%, n=4/45), and AAV2 (7%, n=9/121). Only 1 donor out of 84 was found positive for an IFNγ-positive response to AAV5 (**Figure 1A**). As expected, due to their strong homology, a coprevalence analysis reveals that 26% of positive donors are also reacting to 2 or more serotypes (**Figure 1B** and **Table 1**). It is noteworthy that all the positive donors to AAV2 are also responding to at least another serotype in contrast to AAV4, AAV8, AAV9 and AAVrh10. All of the AAV2-positive donors are positive for an anti-AAV8 cellular response. The only AAV5-positive donor shows a response to AAV8, AAV9 and AAVrh10 as well. This donor has a positive response towards the AAV pool 2 of AAV5, 8, 9 and rh10 suggesting a shared epitope (**Supplementary Table S1**). Regarding the AAV8, 31% of the positive donors are also responding to AAV2 and 54% are also positive to AAV9. Interestingly, 73% of AAV9-positive donors are reactive to this serotype only and 27% were found positive for at least another serotype. We also analyzed the cellular reactivity to AAV over ages. We did not observe a higher cellular reactivity to a given serotype in a specific age-ranged population for any serotype (**Supplementary Table S2**).

**Figure 1:**
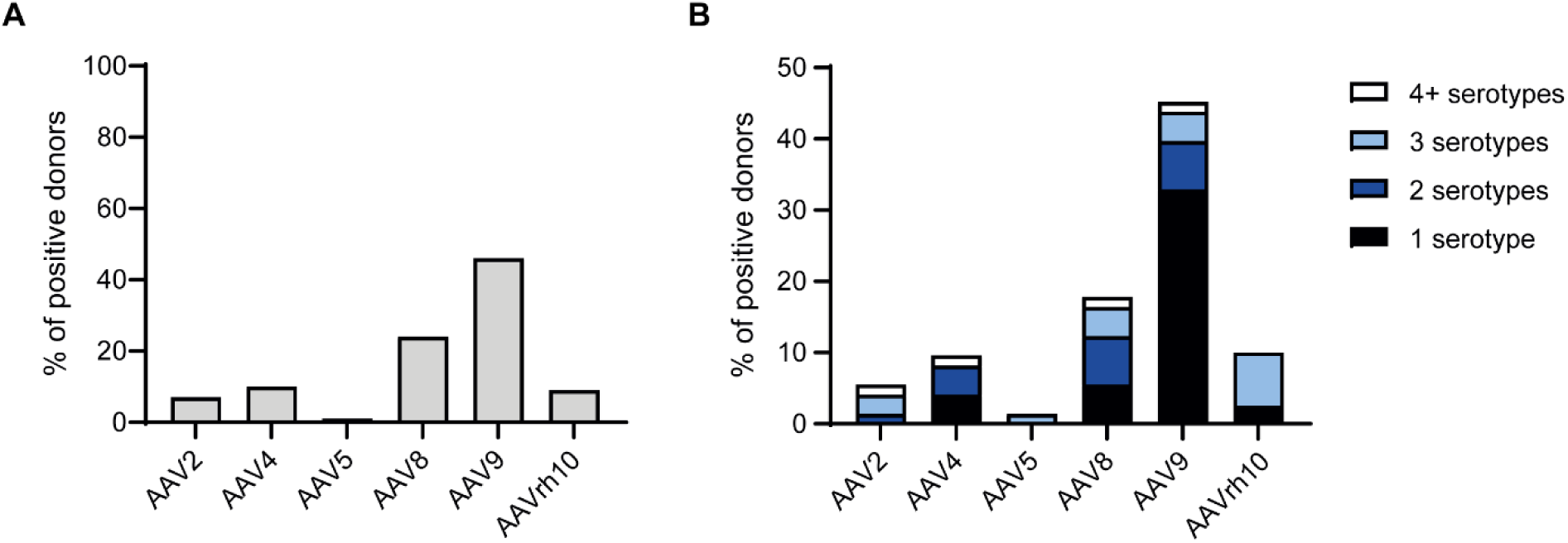
Cellular immune response prevalence to AAV2, 4, 5, 8, 9 and rh10 in healthy population. PBMCs were restimulated *in vitro* with an overlapping peptide library spanning the VP1 sequence of each serotypes tested (*i.e.* AAV2, 4, 5, 8, 9 and Rh10) and analyzed using an IFNγ-ELISpot assay. **A)** Percentage of donors showing an IFNγ-positive response to AAV2 (n=121), AAV4 (n=80), AAV5 (n=84), AAV8 (n=145), AAV9 (n=145), AAVrh10 (n=45). **B)** Coprevalence of the anti-AAV cellular response represented by the percentage of donors showing an IFNγ-positive response to 1, 2, 3 or more than 4 (4+) AAV serotypes. The analysis was performed on 73 healthy donors for AAV2, 4, 5, 8 and 9 and on 40 donors for AAVrh10.

**Table 1:** Coprevalence of cellular immune response to AAV serotypes 2, 4, 5, 8, 9 and rh10. *Anti-AAV positive donors: AAV2: n=4; AAV4: n=7; AAV5, n=1; AAV8: n=13 AAV9: n=33 AAVrh10: n=4*.

|  | AAV2 | AAV4 | AAV5 | AAV8 | AAV9 | AAVrh10 |
| --- | --- | --- | --- | --- | --- | --- |
| AAV2 |  | 25% | 0% | 100% | 75% | 0% |
| AAV4 | 25% |  | 0% | 29% | 43% | N/A |
| AAV5 | 0% | 0% |  | 100% | 100% | 100% |
| AAV8 | 31% | 15% | 8% |  | 54% | 38% |
| AAV9 | 9% | 9% | 3% | 21% |  | 14% |
| AAVrh10 | 0% | N/A | N/A | 75% | 75% |  |

### Cellular and humoral immune responses to AAV2, 8, 9 and rh10 are not correlated

We focused on AAV2 and AAV8 since these serotypes have been studied previously and on AAV9 and rh10 serotypes as they are frequently used in preclinical studies and clinical trials currently and for which there is less data available in the literature. For these 4 serotypes, we investigated the humoral response prevalence by analyzing the presence of anti-AAV total immunoglobulins G (IgGs) by ELISA in the serum of healthy donors (**Figure 2A**). As already reported, we observed a high seroprevalence to AAV2 (40%, n=96, IgG titers from 1:40 to ≥1:10 240, **Supplementary Figure S1A**) and AAV8 (43%, n=109, IgG titers from 1:20 to 1:20 480, **Supplementary Figure S1B**).^5,11^ AAV9 and AAVrh10 show a lower seroprevalence: 29% (n=88, IgG titers from 1:40 to ≥1:10 240, **Supplementary Figure S1C**) and 26% (n=39, IgG titers from 1:640 to ≥1:10 240, **Figure Supplementary S1D**) respectively. We analyzed the coprevalence of AAV-specific IgG in 38 healthy donors (**Table 2**). Eight donors (21%) were positive for the 4 serotypes. Our results show that 85% of the donors positive for anti-AAV2 IgGs are positive for AAV8 and 77% for AAV9 and AAVrh10 capsids. However, in donors positive for anti-AAV8 and anti-AAV9 IgGs, 71 to 79% are positive to AAV2 capsid. 93% of the donors with anti-AAV8 IgGs are responsive to AAV9 and reciprocally. Interestingly, all donors with anti-AAVrh10 antibodies are responding to AAV2 and AAV8 and 90% are responsive to AAV9. The analysis of the humoral reactivity to AAV over ages suggests a higher reactivity to AAV2 and AAVrh10 over time: 23% of the 18-35 yrs old donors (n=31) was positive against 60% for the donors over 55 yrs old (n=25) for the AAV2 and 9% of the 18-35 yrs old donors (n=11) was positive against 50% for the donors over 55 yrs old (n=12) for the AAVrh10 serotype (**Supplementary Table S3**).

**Figure 2:**
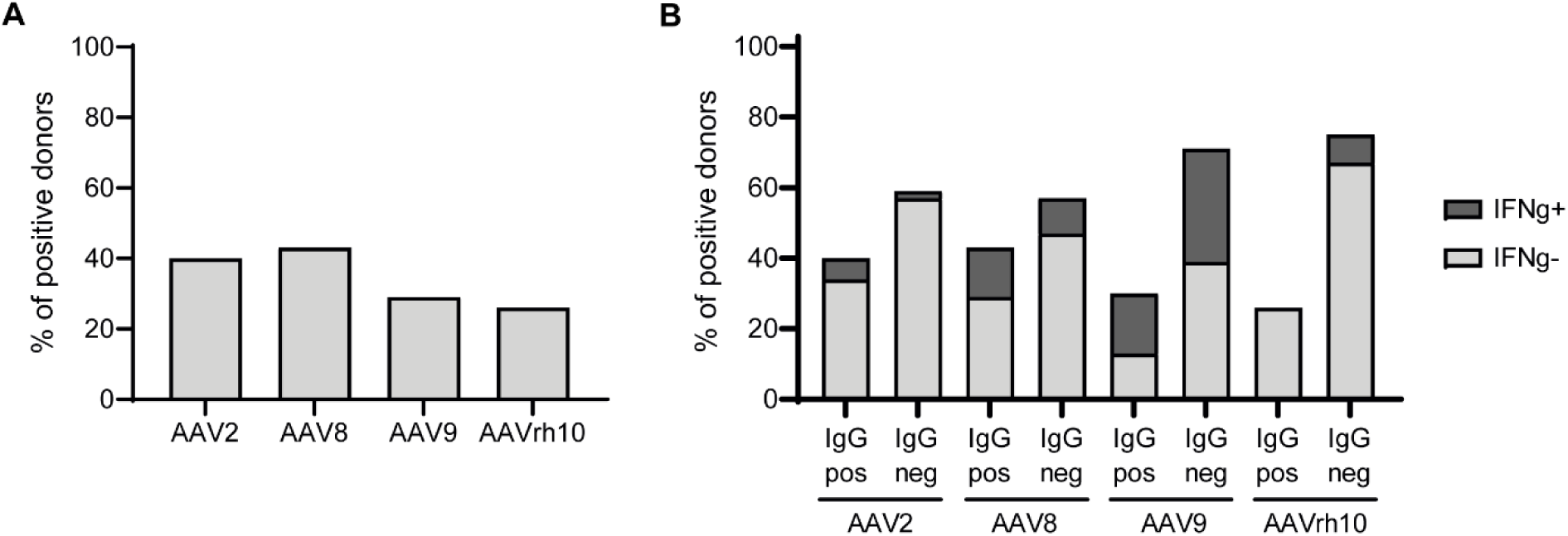
Humoral immune response prevalence to AAV2, 8, 9 and rh10 in healthy population. Sera were collected from healthy human donors and analyzed by ELISA for anti-AAV IgGs. **A)** Percentage of donors showing an anti-AAV IgG response to AAV2 (n=96), AAV8 (n=109), AAV9 (n=92), AAVrh10 (n=39). **B)** Correlation between the anti-AAV humoral and cellular responses in healthy population. Histograms represent percentage of donors with an IFNγ-positive (IFNγ+) or - negative (IFNγ-) response within the healthy population showing an anti-AAV IgG-positive (IgG pos) or -negative (IgG neg) response.

**Table 2:** Coprevalence of humoral immune response to AAV serotypes 2, 4, 5, 8, 9 and rh10. Anti-AAV IgG positive donors: AAV2: n=13; AAV8: n=14 AAV9: n=14 AAVrh10: n=10

|  | AAV2 | AAV8 | AAV9 | AAVrh10 |
| --- | --- | --- | --- | --- |
| AAV2 |  | 85% | 77% | 77% |
| AAV8 | 79% |  | 93% | 71% |
| AAV9 | 71% | 93% |  | 64% |
| AAVrh10 | 100% | 100% | 90% |  |

Finally, we analyzed the correlation between the T cell responses and the antibody responses to AAV in our cohort (**Figure 2B**). For AAV2, in the donors positive for anti-AAV2 IgGs (n=39/96), we observed that 85% of the IgG-positive donors (n=33/39) do not show an IFNγ secretion to the capsid. For AAV8, we observed that 68% of the IgG-positive donors (n=32/47) do not show an IFNγ secretion to the capsid. For AAV9, we observed that 42% of the IgG-positive donors (n=11/26) do not show an IFNγ secretion to the capsid. Surprisingly, none of donors that are positive for anti-AAVrh10 IgG-positive donors (n=0/10) show an IFNγ secretion to the capsid; the donors with a cellular immune response represent 10% of the donors (n=3/29) who do not have anti-AAVrh10 IgGs. Our results confirmed, as previously reported by others, that there is no correlation between the T cell responses and the antibody responses to AAV in our cohort.

### IFNγ-secretion to AAV8 and AAV9 is predominantly mediated by CD8 T cells

Since AAV8 and AAV9 show the highest cellular prevalence (24% and 46% respectively) and are serotypes currently preferred for high systemic dosing in patients, we wanted to further characterize the cellular immune response. We wondered whether CD8 or CD4 T cells were mediating IFNγ secretion. We performed additional IFNγ ELISpot assays in a representative cohort of positive donors (n=8 per serotype) using total PBMCs, CD4- or CD8-depleted PBMCs. The efficiency of each CD4 and CD8 depletion was evaluated by flow cytometry and shows less than 1% of remaining targeted cells after depletion (**Figure 3A** shows one representative donor). These depletions showed that in all positive donors to AAV8 (n=8), IFNγ secretion was mediated by CD8 T cells (i.e. by CD4-depleted PBMCs, **Figure 3B** for one representative donor). No IFNγ secretion was observed in the CD8-depleted fraction for all donors tested (**Supplementary Table S4**). Regarding the AAV9 serotype, 7 positive donors were analyzed and also showed that IFNγ secretion was mediated by CD8 T cells (i.e. by CD4-depleted PBMCs, **Figure 3B**). Nevertheless, in 3 out of 7 donors, in the CD8-depleted PBMCs (*i.e*. CD4 T cells) we also observed a slight IFNγ secretion signal above the negative threshold (dotted line) despite an efficient CD8 depletion (**Figure 3C** for one representative donor and **Supplementary Table S4**). Therefore, IFNγ secretion seems to be mainly mediated by CD8 T cells in response to AAV9 with some donors also showing an IFNγ secretion mediated by CD4 T cells: the depletion of CD4^+^ T cells prior to ELISpot assay noticeably increased the intensity of responses. This is expected since CD4^+^ T cells are removed, CD8^+^ T cells are proportionally more frequent in the tested condition. These results suggest that CD4-depleted cells are the major contributor of IFNγ secretion.

**Figure 3.**
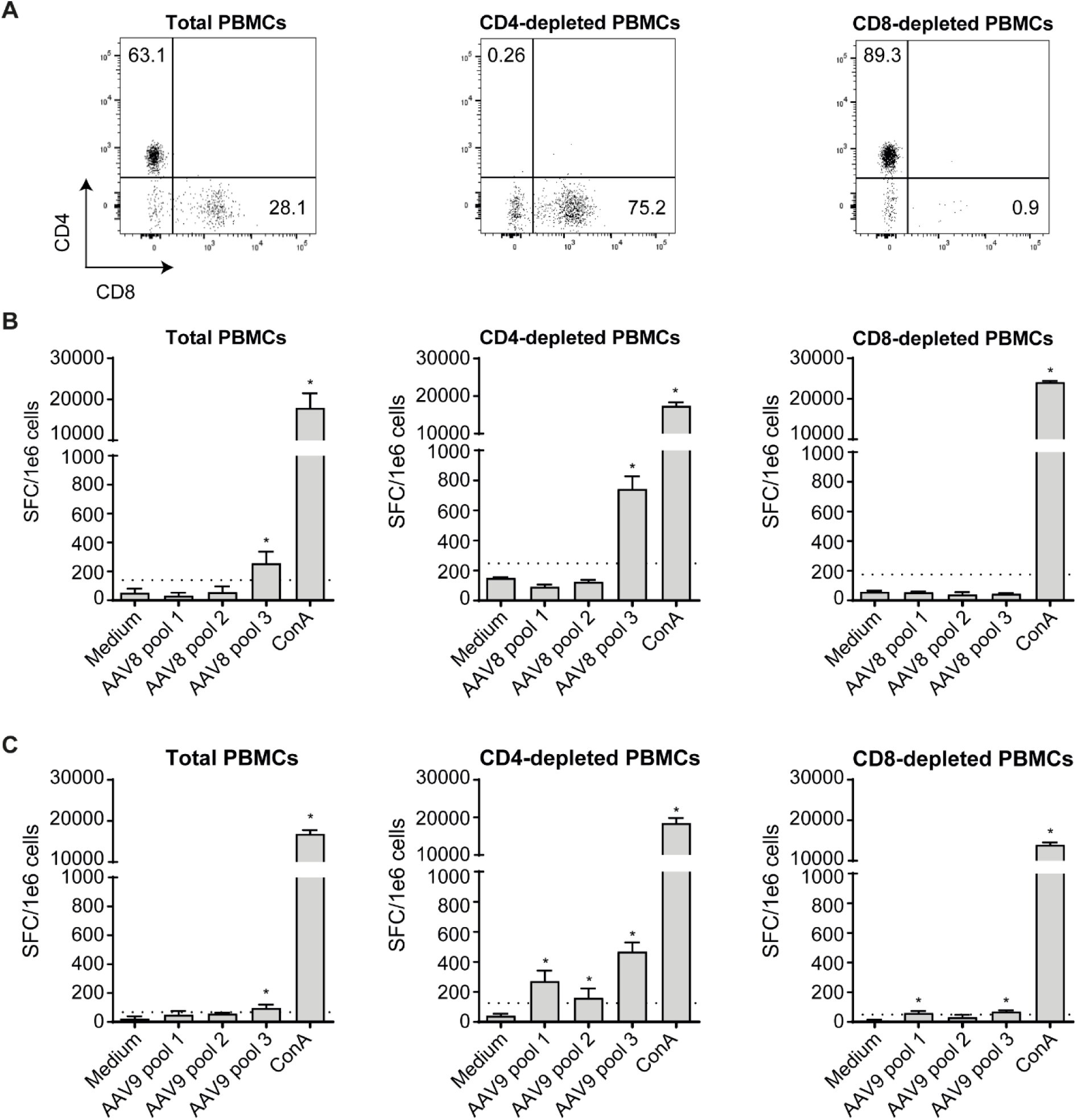
IFNγ secretion to AAV8 and AAV9 is mainly mediated by CD8 T cells. **A)** Cell depletion efficiency analyzed by flow cytometry. Cells were gated from the CD3+CD19-population. Percentages of CD4 (CD4+) and CD8 (CD8+) T cells were determined by flow cytometry before (total PBMCs) and after magnetic separation (CD4- and CD8-depleted PBMCs). (**B**) Total and CD4- and CD8-depleted PBMCs were stimulated *in vitro* with overlapping peptides (15 mers overlapping by 10 aas) spanning the AAV8 VP1 sequence (pool 1–3). This donor is representative of the cohort (n=8). (**C**) Total and CD4- and CD8-depleted PBMCs were stimulated *in vitro* with overlapping peptides (15 mers overlapping by 10 aas) spanning the AAV9 VP1 sequence (pool 1–3). This donor is representative of the cohort showing an IFNγ secretion mediated by both CD4 and CD8 T cells (n=3/7). The 4 remaining donors showed an IFNγ secretion mediated by CD8 T cells only. Negative control consisted of unstimulated cells (medium alone), whereas Concanavalin A (ConA) stimulation was used as a positive control for cytokine secretion. Responses were considered positive when the number of spot-forming colonies (SFCs) per 1e6 cells was >50 and at least 3-fold higher than the negative control condition (dotted line); asterisk (*) denotes a positive response. Error is represented as standard deviation (SD).

### Cytokine secretion profile of preexisting anti-AAV T cells differ between AAV8 and AAV9

We further characterized the preexisting cellular immune response to AAV8 and AAV9 by performing a polyfunctional FluoroSpot assay. We evaluated the secretion of 3 cytokines indicative of a cytotoxic phenotype simultaneously: IFNγ, IL-2 and TNFα (**Figure 4A**). The PBMCs were restimulated *in vitro* with overlapping peptide libraries spanning the AAV8 or AAV9 VP1 sequence. Six and 8 donors showing an IFNγ-positive response to AAV8 and AAV9 respectively were tested (**Figure 4B**). For each donor, we determined within the IFNγ-secreting cell population whether the cells had 1 (IFNγ+ only), 2 (IFNγ+ only and IFNγ+IL-2+ or IFNγ+ only and IFNγ+TNFα+) or 4 functions: (IFNγ+ only, IFNγ +IL-2+, IFNγ+TNFα+ and IFNγ+IL-2+TNFα+). For the AAV8, we observed a diversity in the immune response profile among the 6 donors: 3/6 donors have cells showing 1, 2 and 4 functions, 2/6 donors have cells with 2 functions (IFNγ+ only and IFNγ+TNFα+) and 1/6 donor has cells with 1 function (IFNγ+ only). Surprisingly, for the AAV9, all the donors (n=8) show the same pattern: 2 functions (IFNγ+ only and IFNγ+TNFα+); no IL-2 secretion was detected in any tested donors. Considering the limited number of healthy donors (n=6-8), our results may suggest that preexisting cellular immune responses differ between AAV8 and AAV9 and need to be confirmed with additional healthy donor samples.

**Figure 4:**
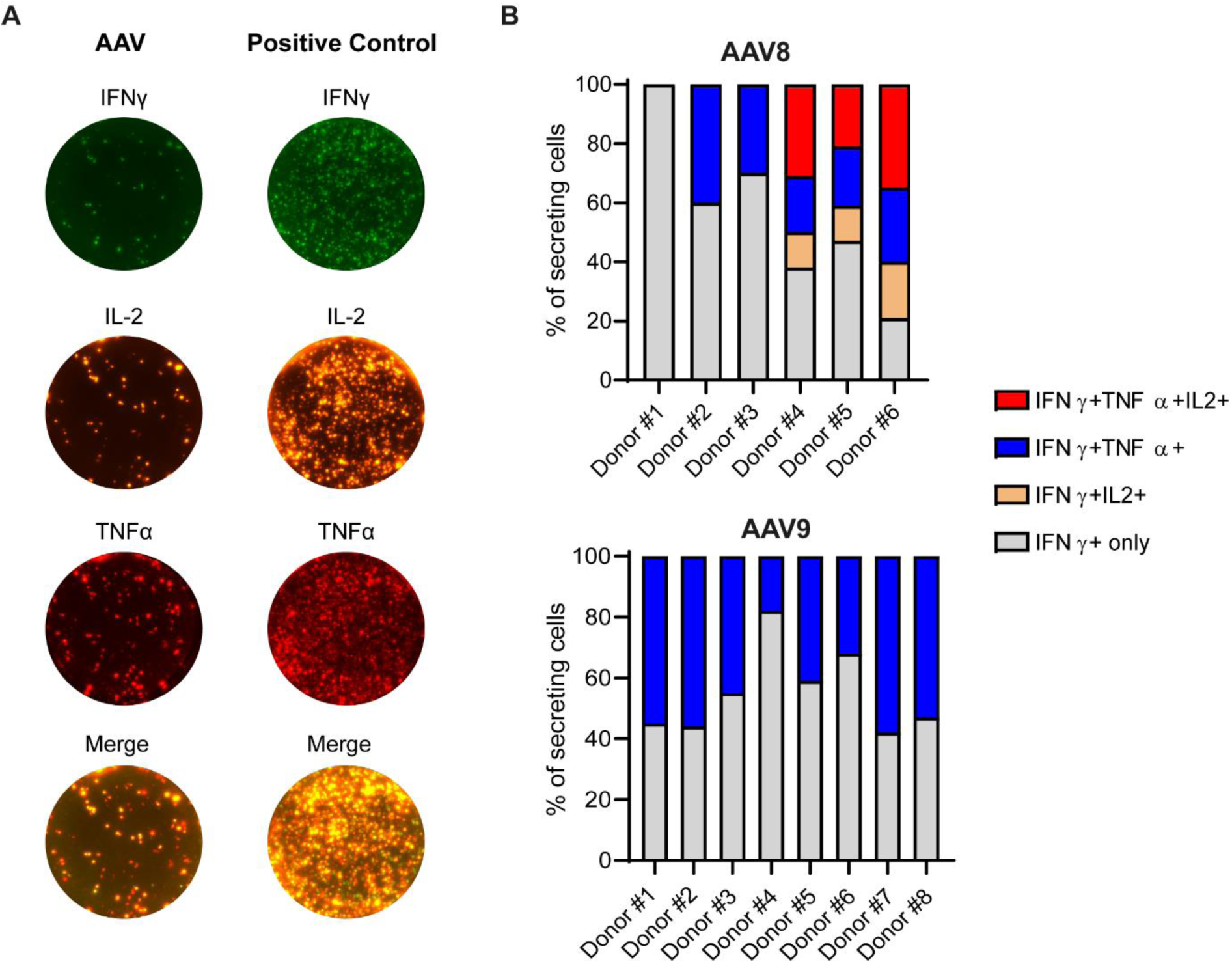
Polyfunctional assessment of anti-AAV8 and anti-AAV9 cellular immune responses. PBMCs were collected from healthy human donors (Donor#1 to #8) and analyzed by IFNγ/IL-2/TNFα FluoroSpot assays. Donors #1 to #6 were previously identified as positive for both anti-AAV8 and anti-AAV9 cellular immune response. Donors #7 and to #8 were identified as positive for anti-AAV9 cellular immune response. **A)** Representative anti-AAV (left) or positive control (right) response using an IFNγ/IL-2/TNFα Flurosopot assay. **B)** Percentage of anti-AAV8 (upper panel, n=6) and anti-AAV 9 (lower panel, n=8) cells within the IFNγ-secreting cell population presenting 1 (IFNγ+ only), 2 (IFNγ+ only and IFNγ+IL-2+ or IFNγ+ only and IFNγ+TNFα+) or 4 functions (IFNγ+ only, IFNγ +IL-2+, IFNγ+TNFα+ and IFNγ+IL-2+TNFα+).

## DISCUSSION

Recombinant AAVs are successful viral vectors for the treatment of inherited disorders as illustrated by recent EMA and FDA approvals of these vectors for the treatment of Leber congenital amaurosis (LCA),^19^ spinal muscular atrophy (SMA),^20^ hemophilia A^21^ and B,^22^ aromatic L-amino acid decarboxylase deficiency (AADC)^23^ and Duchenne muscular dystrophy (DMD)^24^ and more than 230 clinical trials ongoing. However, when protocols were translated from preclinical models to patients, some limits related to the host immune response emerged and remain a major hurdle since protocols evolved from local to systemic injection of high doses of AAV vectors.

Pre-existing immunity to AAV capsid is one of the major limits for successful clinical translation of rAAV gene therapy products. To our knowledge, only the prevalence of pre-existing cellular immunity to AAV1, AAV2, AAV5 and AAV8 has been assessed in the general populations so far.^11–13,25^ Cellular immunity towards more recently emerging serotypes such as AAV9 or AAVrh10 have not been reported yet whereas they are used in 33 and 10 active or recruiting clinical trials respectively (clinicaltrials.gov database). Here, we studied the prevalence of the cellular immune response to 6 AAV serotypes: AAV2, 4, 5, 8, 9 and rh10 in a large cohort of healthy donors. Our results show that AAV9 is most prevalent (46%) followed by AAV8 (24%), AAV4 (10%), AAVrh10 (9%), AAV2 (7%) and AAV5 (1%). The AAV2 prevalence is comparable to results obtained by Chirmule *et al.*^25^ in a cohort of 45 healthy donors from USA but lower than reported by Li *et al.* (47%, n=17)^13^ or by Kruzik *et al.* (19%, n=90).^11^ In this study, they also described a prevalence of 24% for the AAV5 whereas we showed only 1 positive donor out of 84. These differences across studies might be explained by the different assays used and the absence of standardization of these assays (e.g. discrepancy for the positive thresholds determined in different ELISpot assays) but also by an analysis performed on donors coming from different countries or continents (France *versus* Germany, Austria and USA) which has an impact on natural AAV infections as previously shown for humoral prevalence.^4^ These differences highlight the need of standardized assays to compare studies and to measure T-cell mediated immune response to AAV prior and post gene therapy in clinic.

The humoral immune response to AAV2, 8, 9 and rh10 has also been analyzed. As already described, the highest IgG prevalence is observed for AAV2^4,5^ and AAV8 followed by AAV9 and rh10. This is in accordance with previous studies performed on anti-AAV neutralizing antibodies (NAbs) and/or total IgG antibodies (TAbs).^4,5,11,26^ It is noteworthy that some variability may apply between studies since there is no standardized assay with a defined cut-off to measure either NAbs or TAbs in healthy donors or patients enrolled in clinical trials (1:10 in the present study *versus* 1:20^11^ or 1:30^5^ for example). Moreover, the seroprevalence may vary depending on the age of the donors and their origin.^3,4,11^. As previously described, our results confirm there is a high cross reactivity between serotypes; more than 88% of donors responding to AAV2 are responding to AAV8 and 93% of the donors responding to AAV8 are responding to AAV9. The more cross-reactive donors are the ones positive for anti-AAVrh10 (n=9/39). This can be explained by phylogenic relatedness.^27,28^ Our study also confirms there is no correlation between humoral and the IFNγ-mediated cellular immune responses to AAV vectors.^11,12^

As expected, we also report a cellular immune response cross-reactivity between serotypes. This analysis was performed on 73 donors for serotypes 2, 4, 5, 8 and 9 and on 40 donors for serotypes 2, 8, 9 and rh10. Our results show that 26% of the positive donors are also reacting to 2 or more serotypes. All the positive donors to AAV2 were also found responsive to at least another serotype in contrast to AAV4, AAV8, AAV9 and AAVrh10. For AAV2 and AAVrh10, pool 2 elicited a majority of positive response whereas for AAV4, AAV8 and AAV9 it is the pool 3 (**Supplementary Table S1**). Surprisingly, 73% of AAV9-positive donors are reactive to this serotype only whereas the AAV9, AAV8 and AAVrh10 VP1 amino acid sequences share more than 80% of homology (**Supplementary Table S5**). For AAV2, 4 out of 6 of the positive donors were responding to Pool 2 or 3 with 2 donors responding to both peptide pools. These 2 peptide pools contain immunodominant peptides that were previously described in HLA-A*0201 and HLA-B*0702restricted donors.^29^ Some of these immunodominant peptides are also present in AAV9 and AAVrh10 pool 3 although might not be the immunodominant peptides for these particular capsids. Indeed, for these 2 serotypes, donors sharing positive IFNγ-ELISpot do not necessarily respond to the same peptide pools: a positive donor for both AAV2 and AAV9 might have a positive response to AAV2 pool 2 and AAV9 pool 3 (not shown). This observation has been made for all the studied serotype except AAV5 where the only positive donor is also positive for AAV8, AAV9 and AAVrh10 pool 2. An in-depth analysis using a matrix approach would be useful to identify immunodominant peptides depending on donor’s HLA genotype and to develop engineered capsids to lower or prevent the adaptive immune response without altering AAV tropism such as described for neutralizing antibody response.^27,30^

Then, we performed an in-depth analysis of the positive response to AAV8 and AAV9 (which are currently preferred for systemic administration and used in patients (NCT02122952; NCT03199469; NCT03368742). They also appear to be the most prevalent serotypes for a cellular immune response in our healthy donor cohort. Our results show that the pre-existing cellular immune response to AAV8 and AAV9 are mainly mediated by CD8 T cells. Regarding the AAV9, some of the donors are also showing an IFNγ secretion mediated by CD4 T cells. This observation has been also described for AAV2^13^ and AAV1^12^ and reactive cells were identified as CD4+ effector memory T cells. Since CD8 and CD4 depletions were performed on total PBMCs to evidence a CD8- or CD4-mediated response, we cannot exclude that IFNγ secretion might also be mediated by other immune cells especially in seronegative donors. Indeed, Kuranda *et al.* described that CD16^bright^CD56^dim^ natural killer (NK) cells are the major contributor of IFNγ secretion in AAV-seronegative healthy donors towards the AAV2 capsid.^31^

Finally, another major concern in AAV-mediated gene transfer is the fact that IFNγ secretion alone, as measured today, is not sufficient to predict the immune outcome of gene transfer. Actually, the IFNγ secretion is not always associated with a loss of transgene expression^32,33^ suggesting that, aside the vector serotype, the dose or the transgene, the anti-AAV immune response after gene transfer might vary depending on the immune patient profile. While we performed the prevalence study using the standard IFNγ ELISpot assay, we also performed an in-depth characterization using a multiparametric IFNγ/IL-2/TNFα Fluorospot assay on a limited cohort of healthy donors (n=6-8). Our results show a diversity in immune response profile among the 6 AAV8-positive donors whereas the 8 donors tested for AAV9 showed the same pattern with cells that are not secreting IL-2. The absence of IL-2 secretion suggests that these cells may not be cytotoxic effector T cells but rather be exhausted or terminally differentiated effector memory (T_EMRA_).^34,35^ Even though these T cell subtypes are reported in chronic viral infections,^36,37^ they have also been described in the context of AAV. CD8 exhausted T cells expressing the PD1 marker have been described after AAV-mediated gene transfer in mice, non-human primates and humans^38–40^ whereas T_EMRA_ cells reactivity to AAV have been described in healthy donors.^41^ Further investigation including phenotyping assays will be necessary *i)* to confirm that T cells are actually exhausted or terminally differentiated in the present study and *ii)* to better characterize and understand preexisting anti-AAV T cells.

## CONCLUSION

Altogether, our study shows a high cellular prevalence to AAV9 and AAV8 in healthy donors based on IFNγ secretion. Since this cytokine cannot be considered as a specific and exclusive marker for deleterious immune responses as shown in clinical trials, it does not appear relevant to prescreen eligible patients to AAV-derived gene therapies yet in opposition to anti-AAV NAbs or TAbs that can prevent cell transduction^42^ or lead to innate immune pathway activation.^43^ Although, the present study highlights the need of better characterizing the pre-existing T cell response to AAV using multiparametric assays and develop relevant *in vitro* and *in vivo* models to test them and foresee the outcome on gene transfer efficacy.

## ACKNOWLEDGEMENTS

The authors acknowledge the Cytocell - Flow Cytometry and FACS core facility (SFR Bonamy, BioCore, Inserm UMS 016, CNRS UAR 3556, Nantes, France) for its technical expertise and help, member of the Scientific Interest Group (GIS) Biogenouest and the Labex IGO program supported by the French National Research Agency (n°ANR-11-LABX-0016-01). We also acknowledge the UMR1089 Vector Core (Vivem) for providing the AAV vectors for ELISA assays screening, the EFS for providing us with buffy coats and Manon Loirat and Marion Grard for their help on performing ELISA and ELISpot assays.

## AUTHOR CONTRIBUTIONS

Conceptualization: R.X, C.V, O.A and G.G; Investigation: R.X, A.A, C.V, N.J, M.D, C.C, J.L.D, A.C, M.G and G.G; Methodology: R.X, A.A, C.V, O.A and G.G; Visualization: R.X, A.A, O.A and G.G; Writing – Original Draft: R.X and G.G; Writing – Review & Editing, R.X, A.A, M.G, O.A and G.G; Supervision, O.A and G.G. Funding acquisition: O.A. and G.G.

