## Supplementary data for "Prevalence study of cellular capsid-specific immune responses to AAV2, 4, 5, 8, 9 and rh10 in healthy donors"

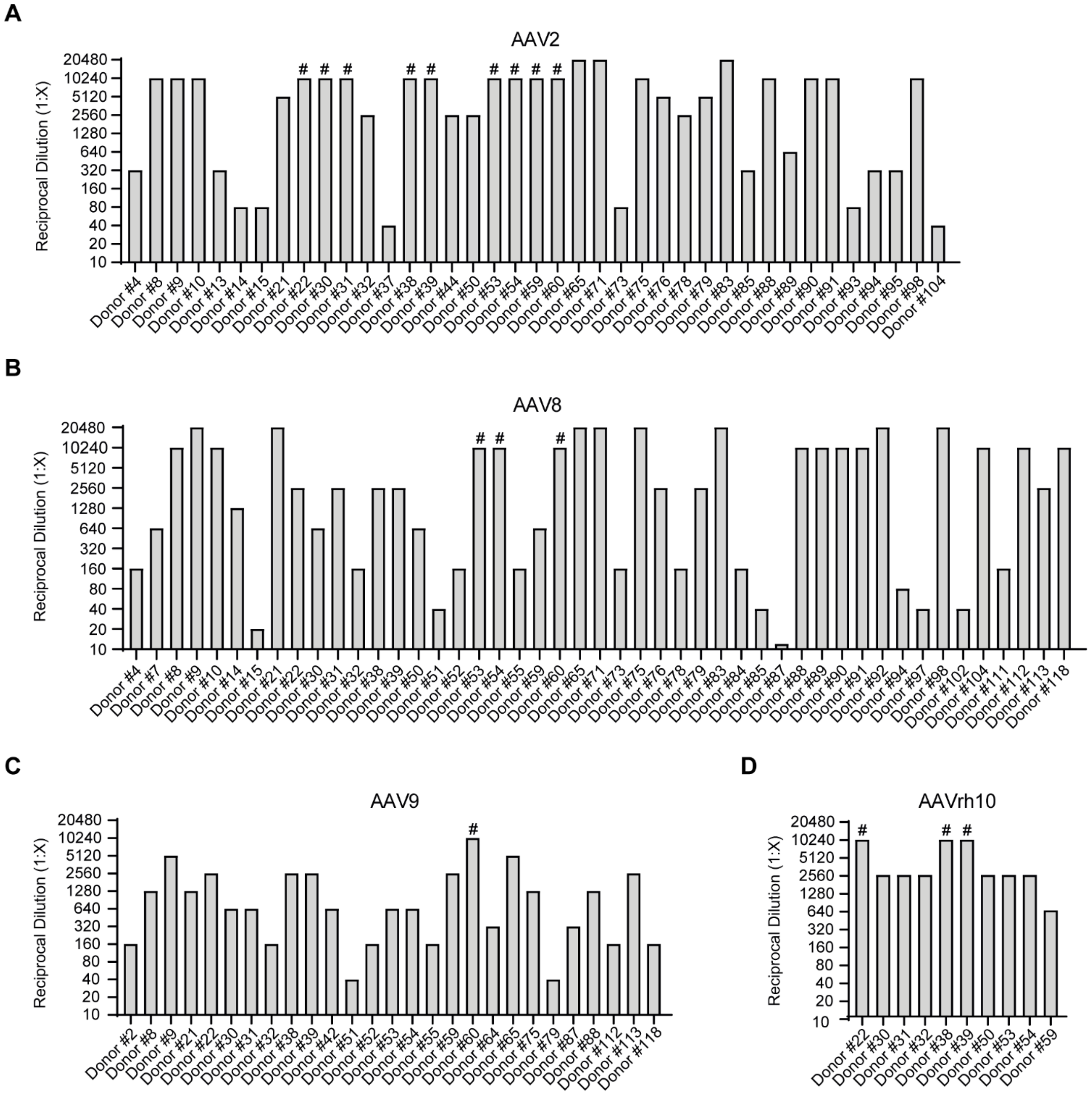


**Figure S1. Anti-AAV IgG titers to AAV2, 8, 9 and rh10 per positive donor.** For each donor, the anti-AAV IgG titer was determined in human plasma by ELISA for serotypes 2 (**A**), 8 (**B**), 9 (**C**) and rh10 (**D**). At least 6 serial 4-fold dilutions starting at 1:10 were performed for each plasma. The titer is defined as the last plasma dilution with an O.D. remaining above the threshold curve.
### indicates the last dilution where the serum was positive for anti-AAV IgGs but the dilution range did not allow to determine the exact titer.

| Serotypes | positive donors/ tested donors | % of positive donors | AAV Pool | positive donors | % of positive donors |
|---|---|---|---|---|---|
| AAV2 | 9/121 | 7% | Pool 1 | 2/9 | 22% |
| | | | Pool 2 | 7/9 | 78% |
| | | | Pool 3 | 4/9 | 44% |
| AAV4 | 8/80 | 10% | Pool 1 | 1/8 | 13% |
| | | | Pool 2 | 3/8 | 38% |
| | | | Pool 3 | 4/8 | 50% |
| AAV5 | 1/84 | 1% | Pool 1 | 0/1 | 0% |
| | | | Pool 2 | 1/1 | 100% |
| | | | Pool 3 | 0/1 | 0% |
| AAV8 | 35/145 | 24% | Pool 1 | 11/35 | 31% |
| | | | Pool 2 | 21/35 | 60% |
| | | | Pool 3 | 24/35 | 69% |
| AAV9 | 66/145 | 46% | Pool 1 | 16/66 | 24% |
| | | | Pool 2 | 25/66 | 38% |
| | | | Pool 3 | 54/66 | 82% |
| AAVrh10 | 4/45 | 9% | Pool 1 | 1/4 | 25% |
| | | | Pool 2 | 3/4 | 75% |
| | | | Pool 3 | 1/4 | 25% |

**Table S1:** Healthy donor reactivity to AAV2, 4, 5, 8, 9 and rh10.

| Age range | AAV2 | AAV4 | AAV5 | AAV8 | AAV9 | AAVrh10 |
|---|---|---|---|---|---|---|
| 18-35 yrs | 11% (n=4/35) | 12% (n=3/25) | 0% (n=0/24) | 26% (n=11/42) | 60% (n=25/42) | 15% (n=2/13) |
| 36-55 yrs | 8% (n=4/48) | 6% (n=2/31) | 0% (n=0/34) | 27% (n=15/56) | 44% (n=25/57) | 6% (n=1/16) |
| >55 yrs | 3% (n=1/35) | 14% (n=3/21) | 4% (n=1/23) | 21% (n=9/43) | 36% (n=15/42) | 6% (n=1/16) |

**Table S2:** Percentage of healthy donors positive for an anti-AAV cellular immune response depending on their ages.

| Age range | AAV2 | AAV8 | AAV9 | AAVrh10 |
|---|---|---|---|---|
| 18-35 yrs | 23% (n=7/31) | 39% (n=13/33) | 25% (n=7/28) | 9% (n=1/11) |
| 36-55 yrs | 42% (n=16/38) | 50% (n=22/44) | 29% (n=11/38) | 19% (n=3/16) |
| >55 yrs | 60% (n=15/25) | 39% (n=12/31) | 36% (n=9/25) | 50% (n=6/12) |

**Table S3:** Percentage of healthy donors positive for anti-AAV IgGs depending on their ages.

| | | IFNγ secretion to AAV | |
|---|---|---|---|
| | | **CD4-depleted** | **CD8-depleted** |
| **AAV8** | Donor #2 | + (Pool 3) | - |
| | Donor #4 | + (Pools 2, 3) | - |
| | Donor #9 | +(Pools 1, 2, 3) | - |
| | Donor #10 | +(Pools 3) | - |
| | Donor #11 | +(Pools 2, 3) | - |
| | Donor #12 | + (Pool 3) | - |
| | Donor #13 | + (Pool 3) | - |
| | Donor #14 | + (Pool 3) | - |
| **AAV9** | Donor #15 | + (Pool 3) | - |
| | Donor #16 | + (Pool 3) | - |
| | Donor #17 | + (Pool 3) | + (Pools 1, 2, 3) |
| | Donor #18 | +(Pools 2, 3) | - |
| | Donor #19 | +(Pools 1, 3) | - |
| | Donor #20 | +(Pools 1, 2, 3) | +(Pool 3) |
| | Donor #21 | +(Pools 1, 2, 3) | +(Pool 3) |

**Table S4:** Cellular immune response to AAV8 and AAV9 mediated by CD4 (*i.e.* CD8-depleted) and/or CD8 (*i.e.* CD4-depleted) T cells.

| | AAV2 | AAV4 | AAV5 | AAV8 | AAV9 | AAVrh10 |
|---|---|---|---|---|---|---|
| **AAV2** | | 61% | 58% | 83% | 82% | 84% |
| **AAV4** | 61% | | 53% | 64% | 63% | 64% |
| **AAV5** | 58% | 53% | | 58% | 57% | 58% |
| **AAV8** | 83% | 64% | 58% | | 85% | 94% |
| **AAV9** | 82% | 63% | 57% | 85% | | 86% |
| **AAVrh10** | 84% | 64% | 58% | 94% | 86% | |

**Table S5:** Percentage of homology of VP1 amino acid sequences across serotypes.
